# Gut eukaryotic communities in pigs: diversity, composition and host genetics contribution

**DOI:** 10.1101/2020.02.18.941856

**Authors:** Yuliaxis Ramayo-Caldas, Francesc Prenafeta, Laura M Zingaretti, Olga Gonzales, Antoni Dalmau, Raquel Quintanilla, Maria Ballester

## Abstract

This study aims to characterize commensal fungi and protists inhabiting the gut of healthy pigs, and explore the putative host genetic control over diversity and composition of pig gut eukaryotes. Fecal fungi and protists communities from 514 Duroc pigs of two sexes and two different ages were characterized by 18S and ITS ribosomal RNA gene sequencing. The gut mycobiota was dominated by yeasts, with a high prevalence of *Kazachstania* spp. Regarding protists, representatives of four genera (*Blastocystis, Neobalantidium, Tetratrichomonas and Trichomitus*) persisted through more than the 80% of the pigs. Heritabilities for the diversity and abundance of gut eukaryotic communities were estimated with the subset of 60 days aged piglets (N=405). Obtained heritabilities ranged from 0.15 to 0.28, indicating a rather limited host-genetic control. A genome wide association study reported genetic variants associated with the fungal α-diversity (SSC6) and with the abundance of *Blastocystis* spp. (SSC6, SSC17 and SSC18). Annotated candidate genes (*IL23R, IL12RB2, PIK3C3, PIK3CD*, *HNF4A* and *TNFRSF9*) were mainly related to immunity, gut homeostasis and metabolic processes. Our results point towards a minor and taxa specific genetic control over the diversity and composition of the pig gut eukaryotic communities.

## Introduction

The gut microbiome harbors thousands of species of archaea, bacteria, viruses and eukaryotes such as protists and fungi that contribute to host biology. The gut eukaryotic communities of monogastric species show a higher interindividual variability, but less abundance, diversity and richness than their bacterial counterparts (Hooks and O’Malley 2019). The so called gut mycobiome of healthy humans is dominated by fungal genera like *Saccharomyces*, *Candida, Penicillim, Aspergillus* and *Malassezia* (Nash et al. 2017), while protist genera such as *Blastocystis, Entamoeba* and *Enteromonas* have been reported in the human gut across different geographical location (Hamad et al. 2016; Laforest-Lapointe and Arrieta 2018; Parfrey et al. 2014). There is currently considerable interest in understanding the mechanisms through which gut commensal eukaryotic communities contribute to host homeostasis and health (Chabé et al. 2017). The role of fungi and protists in the human gut remains poorly understood, but some authors suggest that commensal species may induce innate immune response in the host (Underhill and Iliev 2014), and they could also have other potential benefits (Laforest-Lapointe and Arrieta 2018; Lukeš et al. 2015; Parfrey et al. 2011). For example, *Blastocystis* spp. and non-pathogenic *Entamoeba spp*. have been associated with a healthy and highly diverse gut microbiome ecosystem (Audebert et al. 2016; Chabé et al. 2017; Tito et al. 2019), while *Tritrichomonas musculis* modulates the intestinal immune system and increases host protection against mucosal bacterial infections in mice (Chudnovskiy et al. 2016).

The few studies about gut eukaryotes conducted in pigs (*Sus scrofa domesticus*) used a limited sample size and mainly focused on isolated members of the eukaryote communities or parasite identification under pathogenic conditions (Arfken et al. 2019; Matsubayashi et al. 2014; Shieban 1971; Summers et al. 2019; White et al. 2019; Wylezich et al. 2019). In swine, *Kazachstania* spp. and members of the *Saccharomycetaceae* family are predominant in the gut mycobiota (Arfken et al. 2019; Shieban 1971; Summers et al. 2019; White et al. 2019), while gut protist community of healthy pigs is dominated by *Blastocystis* spp., *Tritrichomonas* spp. and *Balantidium coli*. Environmental factors such as diet seems to play a key role in modulating the structure and composition of both eukaryote and prokaryote communities. However, little is known about the genetic control of the gut eukaryotes, since published studies have been focused on the role of host genetics in shaping the gut bacterial communities (Camarinha-Silva et al. 2017; Chen et al. 2018; Crespo-Piazuelo et al. 2019).

The main goal of this study is to characterize commensal fungi and protists inhabiting the gut of healthy pigs, and explore the putative host genetic control over diversity and composition of gut eukaryotes communities.

## Material and Methods

### Sample collection, DNA extraction and sequencing

Fecal samples were collected at two ages from 514 healthy Duroc pigs belonging to the same commercial outbred line but allocated in two different farms. A total of 405 weaned piglets (204 males and 201 females) distributed in seven batches were sampled in a commercial farm at 60 ± 8 days of age, after four weeks receiving the same transition-based diet. The remaining 109 pigs (50 castrated males and 59 females) were raised under intensive standard conditions at IRTA experimental farm (IRTA, Monells, Spain), and fecal samples were collected at 190 ± 5 days of age, when they fed a finishing standard diet. Both groups of pigs were genetically connected. Animal care and experimental procedures were carried out following national and institutional guidelines for the Good Experimental Practices and were approved by the IRTA Ethical Committee.

DNA was extracted with the DNeasy PowerSoil Kit (QIAGEN, Hilden, Germany), following manufacturer’s instructions. Extracted DNA was sent to the University of Illinois Keck Center for Fluidigm sample preparation and sequencing. The Internal Transcribed Spacer 2 (ITS2) region was amplified using primers ITS3: 5’-GCATCGATGAAGAACGCAGC-3’ and ITS4: 5’-TCCTCCGCTTATTGATATGC-3’ (White et al. 1990). Protist-specific primers (Hadziavdic et al. 2014), F-566: 5’-CAGCAGCCGCGGTAATTCC-3’ and R-1200: 5’-CCCGTGTTGAGTCAAATTAAGC-3’ were used to amplify the 18S rRNA gene fragment. Amplicons were paired-end (2 × 250 nt) sequenced on an Illumina NovaSeq (Illumina, San Diego, CA, USA).

### Bioinformatics and statistical analysis

Sequences were analysed with *Qiime2* (Bolyen et al. 2019), barcode sequences, primers and low-quality reads (Phred scores of <30) were removed. The quality control also trimmed sequences based on expected amplicon length and removed chimeras. Afterwards, sequences were processed into Amplicon Sequences Variants (ASVs) at 99% of identity. ASVs present in less than three samples and representing less than 0.005% of the total counts were filtered out. Samples with less than 6,000 (fungi, n=21 samples) or 10,000 (protists) reads were also excluded. ASVs were classified to the lowest possible taxonomic level based on SILVA v123 database (Quast et al. 2013) for 18S rRNA genes, and the UNITE QIIME version (release 18.11.2018) for fungi (Community 2019.). Subsequently, we excluded those ASVs not taxonomically classified as protists or fungi. Moreover, we verified the fungi taxonomic assignation following the recommendation of Nilsson (Nilsson et al. 2019) by a manual examination of the most abundant fungal ASVs against the International Nucleotide Sequence Database (http://www.insdc.org/). Before the estimation of diversity indexes, samples were rarefied at 6,000 (fungi) and 10,000 (protists) reads of depth, to allow an equal depth. Diversity metrics were estimated with vegan R package (Jari Oksanen et al. 2018). Alpha-diversity was evaluated with the Shannon index (Shannon 1984), and Beta-diversity was assessed using the Whittaker index (Whittaker 1972). To identify environmental (farm, batch or pen) and host-covariates (sex, age, body weight) that may modulate the diversity, structure and profile of the eukaryote communities, we run a Permutational multivariate analysis of variance (PERMANOVA) test using the *adonis* function from vegan (Jari Oksanen et al. 2018). Significance levels were determined after 10,000 permutations and the multiple comparison tests were performed using False Discovery Rate (FDR).

### Estimation of heritability

The putative host genetics determinism of gut eukaryotic profile in pigs was initially assessed by estimating the heritability (*h*^2^) of both fungi and protists alpha-diversity, as well as of their taxa abundance. For these analyses, we used the 405 samples taken at 60 ± 8 days of age in the commercial farm. Parameters estimation was performed using the following Bayesian mixed model implemented with the *bglr* R package (Pérez and de los Campos 2014):

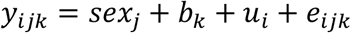

where *y_ijk_* corresponds to the alpha-diversity (fungi or protist) or taxa (genera or specie) clr-transformed abundance of the *i*_th_ individual of sex *j* in the *k*_th_ batch; *sex_j_* and *b_k_* correspond to the systematic effects of *j*_th_ sex (2 levels) and *k*_th_ batch effect (7 levels), respectively; *u*_i_ is the random genetic effect of individual *i*, distributed as 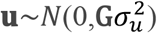, where **G** is the genomic relationship matrix calculated using the filtered autosomal SNPs, finally, *e*_*ijk*_ is the random residual term, with a distribution 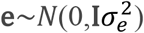. The model was run using a Gibbs sampler with 30,000 iterations and a burn-in of 2,000 rounds. Posterior sample mean and standard deviation of the heritability 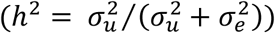 were obtained from the resulted posterior distributions.

### Genotype data and genome wide association study (GWAS)

The Porcine 70k GGP Porcine HD Array (Illumina, San Diego, CA) was used to genotype 390 out of 405 animals sampled at young age in the commercial farm. Quality control was performed with plink (Purcell et al. 2007) to exclude single nucleotide polymorphisms (SNPs) with minor allele frequencies <5%, rates of missing genotypes above 10%, as well as SNPs that did not map to the porcine reference genome (Sscrofa11.1 assembly). Then, to identify SNPs from the host genome associated with the alpha-diversity as well as protists and fungi relative abundances, genome-wide association studies (GWAS) were performed between 42,608 SNPs and the alpha-diversity or the centered log ratio (clr) transformed genera and species abundance tables. Only the genera and species fully taxonomically classified and present in more than 80% of the samples were included in the analysis. For this propose, the genome-wide complex trait analysis (GCTA) software (Yang et al. 2011) was employed using the following model at each SNP:

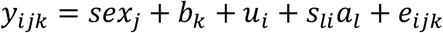

where *y*_*ijk*_ corresponds to the alpha-diversity (fungi or protist) or taxa (genera or specie) clr-transformed abundance of the *i*_th_ individual of sex *j* in the k_th_ batch; *sex_j_*, *b*_*k*_ and *u*_*i*_ are, respectively, the effects of sex, batch and infinitesimal genetic effect described in the previous model, but in this case 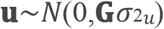 being **G** the genomic relationship matrix calculated using the filtered autosomal SNPs based on the methodology of Yang et al. (Yang et al. 2011); finally, *sli* is the genotype (coded as 0,1,2) for the *l*_th_ SNP of individual *i*, and *al* is the allele substitution effect of SNP *l* on the analysed trait. A SNP was considered to be significantly associated if the corresponding p-value was lower than 0.05, after Benjamini–Hochberg (Benjamini and Hochberg 1995) correction for multiple testing at chromosome level.

### Gene functional classification and canonical pathway analyses

Functional classification and pathway analyses of the annotated candidate genes were carried out using the Ingenuity Pathways Analysis software (IPA; Ingenuity Systems, http://www.ingenuity.com). Significance levels for enrichment of each canonical pathway in the list of candidate genes were calculated using Fisher’s exact test, and the resulting p-values were corrected for multiple-test using the Benjamini and Hochberg algorithm (Benjamini and Hochberg 1995). The cut-off for considering an enrichment as significant was established at a corrected p-value < 0.05.

## Results

### Composition of the pig gut microbial eukaryotic communities

The pig gut microbial eukaryote communities from 514 healthy pigs were analyzed by sequencing the 18S rRNA gene and the ITS2 region. After quality control, 492 fungal and 227 protist ASVs were identified. The pig gut mycobiota was dominated by yeast from the *Kazachstania* genus (average abundance 84%), in particular by the species *K. slooffiae* and *K. bovina* (up to 75.94% and 6.36% abundance, respectively) (**Supplementary table 1**). Other yeasts associated to the *Candida* genus were identified, such as *C. glabrata*, *C. albicans* and *C. (Diutinia) catenulata*, which were present in 1% – 7% of the animals. Besides, a moderate prevalence (15% of pigs) of *Tilletia puccinelliae* was also observed. Along with the ascomycete *Saccharomyces arboricola*, a number of basidiomycetous yeasts were also detected with relatively low incidence (<2%), such as *Sporobolomyces roseus*, *Trichosporon dohaense*, *Debaryomyces prosopidis*, *Pichia sporocuriosa*, *Filobasidium globisporum* and *Vishniacozyma victoriae*. Finally, the pig intestinal mycobiota included a third and more diverse group of cosmopolitan fungal species that are generally categorized as “soil fungi”. In most cases they have a relatively low occurrence in gut (<2%) and are primarily associated with a general saprotrophic ecophisiology (i.e. *Aspergillus* spp., *Aureobasidium pullulans*, *Cladosporium tenuissimum*, *Mucor circinelloides* and *Penicillium polonicum*).

With regard to protists, the prevalent ASV corresponded to an unidentified species of the subclass *Trichostomatia* (superphylum *Alveolata* in the SAR supergroup). Furthermore, the species *Neobalantidium coli* from this class was found in 82% of the animals (**Supplementary table 1)**. Likewise with a relatively high incidence and average abundance (up to 99% and 29%, respectively), a number of *Blastocystis* spp. (superphylum *Stramenopiles* in the SAR supergroup). Other detected commensal trichomonads included *Tetratrichomonas buttreyi* (94% incidence), *Hypotrichomonas imitans* (67% incidence), as well as the *Tetratrichomonas* strain PEKPR (10% incidence), and with a very low incidence and abundance (<1%), *Tritrichomonas suis*. Finally, one representative of the ameboid protist species *Entamoeba gingivalis* (supergroup *Amoebozoa*) was also detected with a relatively high incidence (77%) but a low abundance (<1%).

### Host and environmental factors modulating the diversity of pig gut microbial eukaryotes

The modulatory effect of host and environmental factors over the diversity and composition of gut eukaryotic communities in pigs was evaluated with different approaches. Results from the first PERMANOVA with the whole dataset indicated that the combination of farm and animal age represented the most significant effect shaping the gut eukaryotic communities (p<0.0001), explaining around 44% and 42% of the total variability of fungal and protist communities, respectively. The same data structure was recovered by the principal coordinates analysis (PCoA), which showed two clusters that match with sample farms origin (**Supplementary figure 1**). The diversity index also revealed important differences between farms/ages. Samples taken at 190 d of age in the experimental farm showed a significantly higher protist (p=0.013) and fungal (p=0.006) alpha-diversity. In contrast, weaned piglets raised in commercial conditions showed higher protist beta-diversity (**Supplementary figure 1**). A second PERMANOVA within farm/age was performed. Results of weaned piglets in the commercial farm indicated an important effect of the batch (p-value<0.001) on alpha-diversity of both fungal and protist communities, whereas the pen effect in the second dataset (experimental farm) seems to affect exclusively protist diversity. Regarding sex effects, gut fungi alpha-diversity differed between sexes in weaned piglets (p-value=0.02), whereas no differences between castrated males and females mycobiota diversity were observed at 190 d of age (Table 1).

**Table 1.** Resume of the permutational multivariate analysis of variance.

| **Dataset** | **Community** | **Bacht R2 (P-value)** | **Sex R2 (P-value)** |
| --- | --- | --- | --- |
| Weaned piglets<br>(n=405)<br>commercial farm | Fungi | 0.12 ($p < 0.001$) | 0.01 ($p = 0.02$) |
| | Protist | 0.07 ($p < 0.001$) | 0.02 ($p = 0.60$) |
| Finishing pigs<br>(n=109)<br>experimental farm | Fungi | 0.09 ($p = 0.35$) | 0.01 ($p = 0.24$) |
| | Protist | 0.12 ($p = 0.03$) | 0.01 ($p = 0.55$) |

Afterwards, we implemented a mixed model to assess the degree of host genetic control over the diversity and composition of gut protist and fungal communities. The estimated heritability of the α-diversity index was low, ranging from 0.161±0.053 to 0.188±0.06 for fungi and protists, respectively (Table 2). In the same way, low heritabilities (0.209±0.064) were observed for the abundance of the most representative fungal genus *Kazachstania*, as well as for the protist between 0.159±0.048 and 0.281±0.101 (Table 2). In fact, only three protist genera abundance (*Blastocystis, Trichomitus* and *Neobalantidium*) showed h2 values above 0.2, being the *Blastocysti* abundance (0.281±0.101) the most heritable genera.

**Table 2.** Mean (standard deviation) of marginal posterior distributions for heritabilities of pig gut eukaryotes microbial traits.

| Community | Level | Trait | Heritability (SD) |
| --- | --- | --- | --- |
| Fungi | α-diversity | Shannon-index | 0.161 (0.053) |
|  | Genus | *Kazachstania* | 0.209 (0.064) |
|  | Species | *Kazachstania slooffiae* | 0.216 (0.071) |
| Protist | Alpha-diversity | Shannon-index | 0.188 (0.060) |
|  | Genus | *Blastocystis* | 0.281 (0.101) |
|  |  | *Trichomitus* | 0.233 (0.077) |
|  |  | *Tetratrichomonas* | 0.159 (0.051) |
|  |  | *Neobalantidium* | 0.273 (0.090) |
|  | Species | *Trichomitus batrachorum* | 0.240 (0.075) |
|  |  | *Tetratrichomonas buttrey* | 0.158 (0.048) |
|  |  | *Neobalantidium coli* | 0.256 (0.086) |
|  |  | *Blastocystis* sp. ATCC 50177 | 0.245 (0.086) |
|  |  | *Blastocystis* sp. CK86-1 | 0.203 (0.066) |
|  |  | *Blastocystis* sp. SY94-3 | 0.205 (0.065) |

### Identification of host genetic regions linked with the composition and diversity of pig gut microbial eukaryotes

Results from GWAS revealed few and weak association signals between the host genome and the gut microbial eukaryotes composition and diversity. We identified a total of 174 SNPs as significantly (at chromosome-wide level, FDR<0.05) related with the abundance of protist community (**Supplementary table 2**), located in seven intervals distributed across three pig *Sus scrofa* chromosomes: SSC6, SSC17 and SSC18 (Table 3). The 32.18% of these SNPs were intronic variants, 56.89% were intergenic, 5.75% were located upstream/downstream of genes, 1.15% were exonic synonymous, and 4% mapped within non-coding transcript variants (**Supplementary table 2**).

**Table 3.** Resume of the chromosomal intervals identified by the GWAS

| Chr | Start (bp) | End (bp) | Size (Mb) | Associated microbial trait |
| --- | --- | --- | --- | --- |
| 6 | 68999679 | 69466937 | 0,467 | *Blastocystis* sp. SY94-3 |
| 6 | 120437893 | 121498044 | 1,060 | *Blastocystis* sp. ATCC 50177;<br>*Blastocystis* sp. CK86-1 |
| 6 | 122778463 | 132592988 | 9,814 | *Blastocystis* sp. ATCC 50177;<br>*Blastocystis* sp. CK86-1 |
| 6 | 141914558 | 145392721 | 3,478 | *Blastocystis* sp. ATCC 50177;<br>*Blastocystis* sp. CK86-1; ITS α-diversity |
| 17 | 46764111 | 46988559 | 0,224 | *Blastocystis* sp. |
| 18 | 4538602 | 4570818 | 0,032 | *Blastocystis* sp. ATCC 50177 |
| 18 | 25850358 | 25885196 | 0,035 | *Blastocystis* sp. ATCC 50177 |

Most of these associated SNPs (164 out of 174) were identified on SSC6 (Table 3) and were mainly associated with the relative abundance of two species of *Blastocystis* genera: *CK86-1* and *ATCC 50177* (**Supplementary table 2**). The remaining significant SNPs located on SSC17 (46.77– 46.99 Mb interval) and SSC18 (two intervals: 4,53-4,57 Mb and 25,85-25,88 Mb) were also associated to the relative abundance of members of *Blastocystis* (Table 3). Regarding diversity, the aforementioned 141.91-145.39 Mb region of SSC6 resulted also associated with the fungi Shannon diversity-index (Figure 1). Finally, no significant associations with either the relative abundance of fungi (*Kazachstania* genera or *Kazachstania slooffiae*) or the diversity of protist were observed.

**Figure 1.**
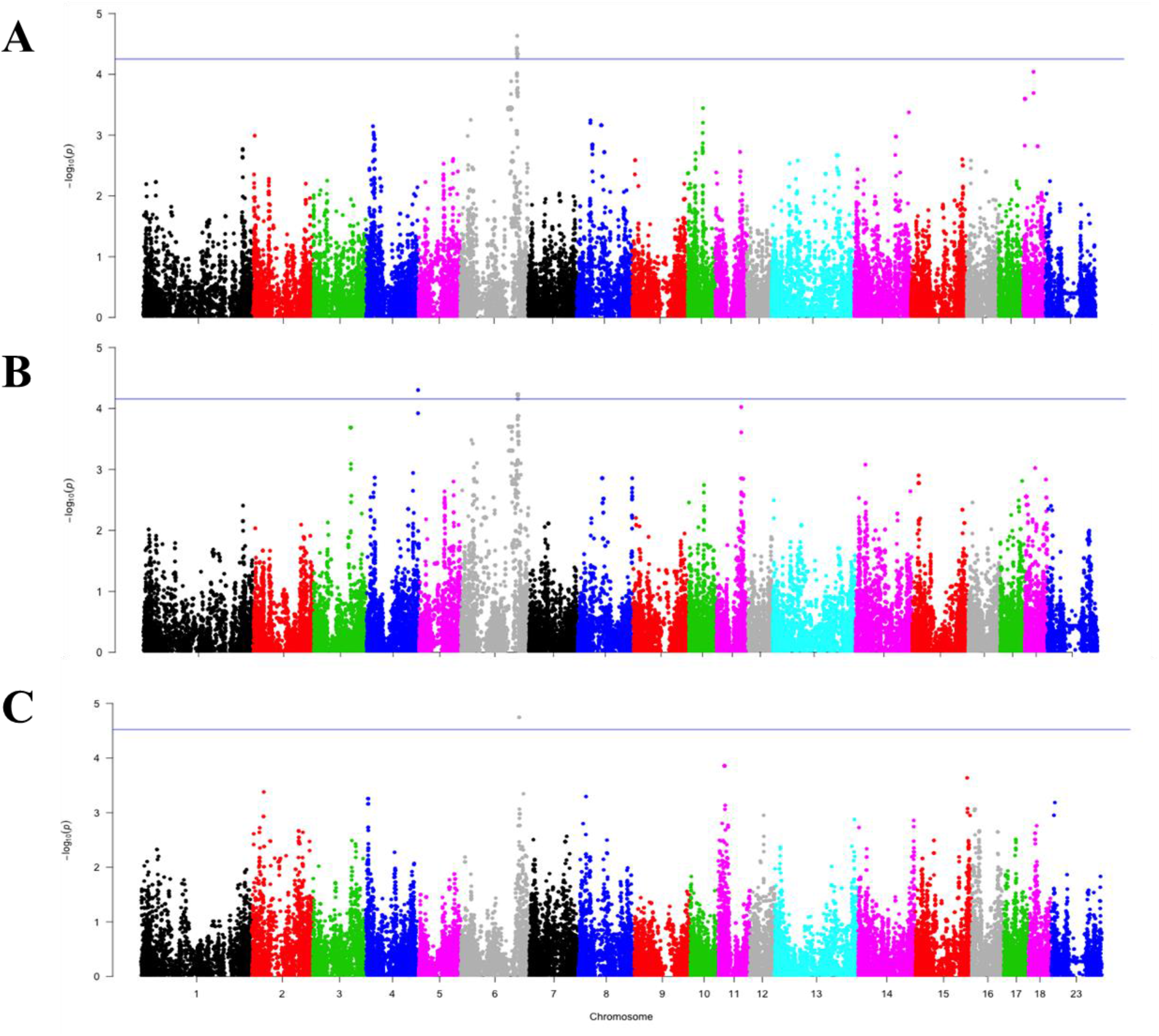
GWAS plot corresponding to A: *Blastocystis* sp. ATCC 50177, B: *Blastocystis* sp. CK8166 and C: Fungi Shannon diversity-index

### Genes and pathways associated to pig gut microbial eukaryotes

A total of 229 protein-coding, 74 long non-coding RNA and two miRNAs (*mir138-2* and *ssc-mir-186*) (**Supplementary table 3**) were annotated within the host genomic intervals associated to the gut microbial eukaryotes composition and diversity. A detailed exploration of annotated genes revealed that the Interleukin 23 Receptor gene (*IL23R*) holds two intronic SNPs, rs343576943 (p-value=1,8*10^−5^) and rs81392455 (p-value= 1,8*10^−5^) that were associated with fungal α-diversity (**Supplementary table 2**). Furthermore, the variants rs81391297, rs81391299 and rs81299019, mapped within the same 122.8-132.6 Mb interval on SSC6 and associated with the relative abundance of *Blastocystis* spp., were located on intron eight of the Phosphatidylinositol 3-Kinase Catalytic Subunit Type 3 (*PIK3C3*) gene. On SSC17, the intronic variant rs331945396 (SSC17:46832698-46832698) in the Hepatocyte Nuclear Factor 4 Alpha (*HNF4A*) gene was also associated with the relative abundance of *Blastocystis* genera (p-value= 9,02*10^−5^). Other genes related to the immune system were annotated within the aforementioned chromosomal intervals, as for example the TNF Receptor Superfamily Member 9 (*TNFRSF9*), the Interleukin 12 Receptor Subunit Beta 2 (*IL12RB2*) or the Phosphatidylinositol-4,5-Bisphosphate 3-Kinase Catalytic Subunit Delta (*PIK3CD*), as well as other members of immune-related pathways (e.g. *CDH1*, *CDH11*, *CDH3*, *CDH5*, *LPAR3*, *PIK3C3*, *CCL17*, *CCL22*, *CDH5*, *CKLF* and *MMP15*).

Finally, the functional annotation revealed that these genes belong to a variety of physiological processes and gene networks related to Cell-To-Cell Signaling and Interaction, Cellular Assembly and Organization, Cellular Function and Maintenance, Developmental Disorder, Hereditary Disorder, Cell-mediated Immune Response, Humoral Immune Response, Immune Cell Trafficking, and Protein Synthesis (**Supplementary table 4**). Metabolic pathways most significantly enriched by the list of candidate genes include Relaxin Signaling, Leptin Signaling in Obesity, IGF-1 Signaling, PXR/RXR Activation and HIF1α Signaling pathways. It should be noted the overrepresentation of immune-related pathways such as Gα12/13 Signaling, IL-9 Signaling, IL-23 Signaling, IL-12 Signaling, ERK/MAPK Signaling, Agranulocyte and Granulocyte Adhesion and Diapedesis (**Supplementary table 5**).

## Discussion

The analysis of gut microbial eukaryotic communities in 514 healthy animals have allowed characterizing commensal fungi and protists inhabiting the porcine gut tract. Our results evidenced that pigs gut mycobiota is dominated by the yeast *Kazachstania* spp. which corresponds to the teleomorphic state of *Candida slooffiae* and *Candida bovina* (Kurtzman et al. 2005). Particularly the species *K. slooffiae* was found in all studied animals across farm, sex and ages, observation that is in agreement with the ubiquitous detection of this yeast from the gut of healthy pigs, and further supports the hypothesis that pigs intestinal track is the primary ecological niche for *K. slooffiae* (Summers et al. 2019). The closely related *K. bovina* was detected in the 45% of the studied pigs, and might also play a significant role in the gut. Several transient fungi that probably arise from the feeding or the environment were observed but with low incidence. The plant-gut association for certain yeast is more evident in a second group of filamentous fungi that are characterized as specialized plant biotrophs. The most representative of those is the basidiomycete *T. puccinelliae* that, surprisingly, occurred in 15% of the studied pigs. Species from *Tilletia* genera are smut fungi that infect various grasses from the *Poaceae* family and encompass plant pathogens of economic importance in the production of cereals and forage grasses. So far, *T. puccinelliae* has only been isolated from weeping alkaligrass (*Puccinellia distans*), a common ruderal grass in Europe and North America (Bao et al. 2010). Plants infected by other *Tilletia* species do not pose a toxicity risk for humans but contaminated grains might be derived to animal feed due to off-flavors (Preugschat et al. 2014). Such association between feed and fungal gut might also apply for the plant pathogens *Ustilago hordei* and *Mycosphaerella tassiana*, found in about 3% of the animals. Former results are in agreement with previous reports on humans and non-human primates (Hallen-Adams et al. 2015; Mann et al. 2019; Nash et al. 2017; Raimondi et al. 2019), and suggest that, differently to pig gut prokaryotes (Xiao et al. 2016), the pig gut mycobiome may lack a stable core. Consequently, it is expected that a large proportion of the fungi detected in pig fecal samples may be transients from dietary or environmental origin.

Meanwhile, the prevalence (>80% of the 514 pig samples) of four protist genera (*Blastocystis, Neobalantidium*, *Tetratrichomonas* and *Trichomitus*) was observed. *Neobalantidium* is a world-wide parasitic-opportunistic human pathogen that is transmitted through the fecal-oral route, particularly when there is a close contact with pigs, which are its asymptomatic reservoir hosts (Schuster and Ramirez-Avila 2008). Similarly, several species in *Blastocystis* are causative agents of diarrhea in humans through the fecal-oral infection route, being the pig a common reservoir (Rivera 2008). A number of protists in the *Trichomonadida* order (supergroup *Excavata*) that are common in the digestive tract of pigs (Mostegl et al. 2012) were also found in the present study. Colonization with this species has not been associated to virulence in mammals, but the full pathogenic potential of *T. batrachorum* has yet to be explored.

The analysis of host intrinsic and extrinsic factors ascertained the significant modulatory role of the environmental effects gathered in batch and farm factors (e.g. climate, management conditions, diet) on gut eukaryotes composition and diversity. Also important differences between gut eukaryotic communities in weaned and finishing pigs were evidenced, whereas scarce differences due to the sex of the animal were observed. Subsequently, we explored the putative host genetic control. The low heritabilities estimates (ranging from 0.16 to 0.28) allowed inferring a limited and taxa specific host genetic control over diversity and composition of pig gut eukaryotes communities. Nevertheless, GWAS revealed genetic variants associated with fungal α-diversity (on SSC6) and the abundance of *Blastocystis* spp (on SSC6, 17 and 18). Interestingly, two intronic SNPs associated with fungal α-diversity were mapped on the *IL23R* gene that had been reported to be involved in the diversity of ileum microbiota in humans (Zakrzewski et al. 2018). In the same chromosome, *PIK3C3* gene gathered two SNPs associated to *Blastocystis* abundance. *PIK3C3* is required for T cell homeostasis (McLeod et al. 2011; Parekh et al. 2013; Willinger and Flavell 2012) and reported to play a relevant role in maintaining gut homeostasis (Zhao et al. 2018). Other genes involved in the intestinal epithelial integrity and gut homeostasis were annotated on SSC17, such as *HNF4A*, a relevant regulator that mediates microbial control of intestinal gene expression (Davison et al. 2017). Also a number of genes related to the immune system were annotated within the aforementioned chromosomal intervals, as the *TNFRSF9* gene that contributes to the development, survival and activation of T cells (Schwarz et al. 1993; Ward-Kavanagh et al. 2016), the *IL12RB2* and *PIK3CD* genes, and a plethora of members of the immune-related pathways Gα12/13 Signaling, IL-9 Signaling, IL-23 Signaling, IL-12 Signaling, ERK/MAPK Signaling, Agranulocyte and Granulocyte Adhesion and Diapedesis.

These findings are in line with recent reports in mice (Khan et al. 2019), and suggest that polymorphisms located in (or in linkage disequilibrium with) genes related to immune system and gut homeostasis may modulate the diversity and composition of the eukaryote gut communities in pigs. The evolutionary mechanism beyond the observed association may be explained by indirect relationships between mutual favorable selection at host-genome and microbial level (Wang et al. 2018; Zilber-Rosenberg and Rosenberg 2008). Although far to be fully understood, some examples of host-genetic microbiota associations have been reported (reviewed in (Wang et al. 2018)), that similar to our results identify candidate genes mainly related to host-response against pathogens, sensing microbes, cell signaling pathways and innate immune system. However, in agreement with previous reports (Khan et al. 2019; Rothschild et al. 2018), ours findings also indicate a limited and target specific host genetic control over the composition of the gut eukaryote communities in pig.

In summary, our findings highlight the relevance of considering the gut eukaryotic communities to better understand the porcine gastrointestinal microbiome ecosystem. We are aware of some limitations of our study, as for example the limited sample size or the lack of degrees of freedom to estimate all effects, but also of methodological constraints such as primers choice or the use of less curated (compared to bacterial) protists and fungal databases, that together with the reduced amount of public available reference genomes compromise the accuracy of the taxonomic classification. To the best of our knowledge, this study represents the largest effort to characterize the gut fungal and commensal protist communities in pigs, but further larger studies including experimental validations and alternative meta-sequencing approaches are needed to unveil the role of the host-associated microbial communities in pigs production performance, welfare and health.

## Conclusions

The diversity and composition of gut commensal eukaryotic communities in healthy pigs at two ages have been characterized. The porcine gut mycobiota is dominated by yeast, with a high prevalence of *Kazachstania*, and a common set of four protist genera (*Blastocystis, Neobalantidium, Tetratrichomonas and Trichomitus*) persisted through the majority of animals. Our results point towards a minor and taxa specific genetic control over the diversity and composition of the pig gut eukaryotic communities, but we describe associations with genes functionally related to the immune system and gut homeostasis that might have an effect in modulating the fungi α-diversity and the abundance of *Blastocystis* ssp.

## Acknowledgements

The authors warmly thank all technical staff from *Selección Batallé* S.A, for providing the animal material and their collaboration during the sampling, as well as to Juan Pablo Sanchez for valuable discussions and comments on the manuscript. Y. Ramayo-Caldas was funded by Marie Skłodowska-Curie grant (P-Sphere) agreement No 6655919 (EU). L.M. Zingaretti is recipient of a Ph.D. grant from Ministry of Economy and Science, Spain associated with ‘Centro de Excelencia Severo Ochoa 2016–2019’ award SEV-2015-0533 to CRAG. M. Ballester is recipient of a Ramon y Cajal post-doctoral fellowship (RYC-2013–12573) from the Spanish Ministry of Economy and Competitiveness. Part of the research presented in this publication was funded by Grants AGL2016-75432-R, AGL2017-88849-R awarded by the Spanish Ministry of Economy and Competitiveness. The authors belong to Consolidated Research Group TERRA (AGAUR, ref. 2017 SGR 1290 and 2017 SGR 1719).

## References

Arfken AM, Frey JF, Ramsay TG, and Summers KL. 2019. Yeasts of Burden: Exploring the Mycobiome–Bacteriome of the Piglet GI Tract. Front Microbiol 10:2286.

Audebert C, Even G, Cian A, Safadi DE, Certad G, Delhaes L, Pereira B, Nourrisson C, Poirier P, Wawrzyniak I et al. . 2016. Colonization with the enteric protozoa Blastocystis is associated with increased diversity of human gut bacterial microbiota. Scientific Reports 6(1):25255.

Bao X, Carris LM, Huang G, Luo J, Liu Y, and Castlebury LA. 2010. Tilletia puccinelliae, a new species of reticulate-spored bunt fungus infecting Puccinellia distans. Mycologia 102(3):613–623.

Benjamini Y, and Hochberg Y. 1995. Controlling the False Discovery Rate: A Practical and Powerful Approach to Multiple Testing. Journal of the Royal Statistical Society Series B (Methodological) 57(1):289–300.

Bolyen E, Rideout JR, Dillon MR, Bokulich NA, Abnet CC, Al-Ghalith GA, Alexander H, Alm EJ, Arumugam M, Asnicar F et al. . 2019. Reproducible, interactive, scalable and extensible microbiome data science using QIIME 2. Nature Biotechnology 37(8):852–857.

Camarinha-Silva A, Maushammer M, Wellmann R, Vital M, Preuss S, and Bennewitz J. 2017. Host Genome Influence on Gut Microbial Composition and Microbial Prediction of Complex Traits in Pigs. Genetics 206(3):1637.

Chabé M, Lokmer A, and Ségurel L. 2017. Gut Protozoa: Friends or Foes of the Human Gut Microbiota? Trends in Parasitology 33(12):925–934.

Chen C, Huang X, Fang S, Yang H, He M, Zhao Y, and Huang L. 2018. Contribution of Host Genetics to the Variation of Microbial Composition of Cecum Lumen and Feces in Pigs. Front Microbiol 9:2626.

Chudnovskiy A, Mortha A, Kana V, Kennard A, Ramirez JD, Rahman A, Remark R, Mogno I, Ng R, Gnjatic S et al. . 2016. Host-Protozoan Interactions Protect from Mucosal Infections through Activation of the Inflammasome. Cell 167(2):444–456.e414.

Community U. 2019. UNITE QIIME release for Fungi. Version 18.11.2018. UNITE Community. https://doi.org/10.15156/BIO/786334.

Crespo-Piazuelo D, Migura-Garcia L, Estellé J, Criado-Mesas L, Revilla M, Castelló A, Muñoz M, García-Casco JM, Fernández AI, Ballester M et al. . 2019. Association between the pig genome and its gut microbiota composition. Scientific Reports 9(1):8791.

Davison JM, Lickwar CR, Song L, Breton G, Crawford GE, and Rawls JF. 2017. Microbiota regulate intestinal epithelial gene expression by suppressing the transcription factor Hepatocyte nuclear factor 4 alpha. Genome Res 27(7):1195–1206.

Hadziavdic K, Lekang K, Lanzen A, Jonassen I, Thompson EM, and Troedsson C. 2014. Characterization of the 18S rRNA Gene for Designing Universal Eukaryote Specific Primers. PLoS One 9(2):e87624.

Hallen-Adams HE, Kachman SD, Kim J, Legge RM, and Martínez I. 2015. Fungi inhabiting the healthy human gastrointestinal tract: a diverse and dynamic community. Fungal Ecology 15:9–17.

Hamad I, Raoult D, and Bittar F. 2016. Repertory of eukaryotes (eukaryome) in the human gastrointestinal tract: taxonomy and detection methods. Parasite Immunology 38(1):12–36.

Hooks KB, and O’Malley MA. 2019. Contrasting Strategies: Human Eukaryotic Versus Bacterial Microbiome Research. Journal of Eukaryotic Microbiology n/a(n/a).

Jari Oksanen F, Blanchet G, Friendly M, Kindt R, Legendre P, McGlinn D, Minchin PR, O’Hara RB, Simpson GL, Solymos P et al. . 2018. vegan: Community Ecology Package. R package version 2.5-3. https://CRANR-projectorg/package=vegan.

Khan AA, Yurkovetskiy L, O’Grady K, Pickard JM, de Pooter R, Antonopoulos DA, Golovkina T, and Chervonsky A. 2019. Polymorphic Immune Mechanisms Regulate Commensal Repertoire. Cell Reports 29(3):541–550.e544.

Kurtzman CP, Robnett CJ, Ward JM, Brayton C, Gorelick P, and Walsh TJ. 2005. Multigene phylogenetic analysis of pathogenic candida species in the Kazachstania (Arxiozyma) telluris complex and description of their ascosporic states as Kazachstania bovina sp. nov., K. heterogenica sp. nov., K. pintolopesii sp. nov., and K. slooffiae sp. nov. J Clin Microbiol 43(1):101–111.

Laforest-Lapointe I, and Arrieta M-C. 2018. Microbial Eukaryotes: a Missing Link in Gut Microbiome Studies. mSystems 3(2):e00201–00217.

Lukeš J, Stensvold CR, Jirků-Pomajbíková K, and Wegener Parfrey L. 2015. Are Human Intestinal Eukaryotes Beneficial or Commensals? PLOS Pathogens 11(8):e1005039.

Mann AE, Mazel F, Lemay MA, Morien E, Billy V, Kowalewski M, Di Fiore A, Link A, Goldberg TL, Tecot S et al. . 2019. Biodiversity of protists and nematodes in the wild nonhuman primate gut. The ISME Journal.

Matsubayashi M, Suzuta F, Terayama Y, Shimojo K, Yui T, Haritani M, and Shibahara T. 2014. Ultrastructural characteristics and molecular identification of Entamoeba suis isolated from pigs with hemorrhagic colitis: implications for pathogenicity. Parasitology Research 113(8):3023–3028.

McLeod IX, Zhou X, Li Q-J, Wang F, and He Y-W. 2011. The Class III Kinase Vps34 Promotes T Lymphocyte Survival through Regulating IL-7Rα Surface Expression. The Journal of Immunology 187(10):5051.

Mostegl MM, Richter B, Nedorost N, Lang C, Maderner A, Dinhopl N, and Weissenböck H. 2012. First evidence of previously undescribed trichomonad species in the intestine of pigs? Veterinary parasitology 185(2-4):86–90.

Nash AK, Auchtung TA, Wong MC, Smith DP, Gesell JR, Ross MC, Stewart CJ, Metcalf GA, Muzny DM, Gibbs RA et al. . 2017. The gut mycobiome of the Human Microbiome Project healthy cohort. Microbiome 5(1):153.

Nilsson RH, Anslan S, Bahram M, Wurzbacher C, Baldrian P, and Tedersoo L. 2019. Mycobiome diversity: high-throughput sequencing and identification of fungi. Nature Reviews Microbiology 17(2):95–109.

Parekh VV, Wu L, Boyd KL, Williams JA, Gaddy JA, Olivares-Villagómez D, Cover TL, Zong W-X, Zhang J, and Van Kaer L. 2013. Impaired autophagy, defective T cell homeostasis, and a wasting syndrome in mice with a T cell-specific deletion of Vps34. J Immunol 190(10):5086–5101.

Parfrey LW, Walters WA, and Knight R. 2011. Microbial eukaryotes in the human microbiome: ecology, evolution, and future directions. Front Microbiol 2:153–153.

Parfrey LW, Walters WA, Lauber CL, Clemente JC, Berg-Lyons D, Teiling C, Kodira C, Mohiuddin M, Brunelle J, Driscoll M et al. . 2014. Communities of microbial eukaryotes in the mammalian gut within the context of environmental eukaryotic diversity. Front Microbiol 5:298–298.

Pérez P, and de los Campos G. 2014. Genome-wide regression and prediction with the BGLR statistical package. Genetics 198(2):483–495.

Preugschat K, Kersten S, Ettle T, Richter W, Karl H, Breves G, Büttner P, and Dänicke S. 2014. Effects of feeding diets containing increasing proportions of bunt-infected wheat (Tilletia caries) on performance and health of pigs. Archives of Animal Nutrition 68(1):55–62.

Purcell S, Neale B, Todd-Brown K, Thomas L, Ferreira MAR, Bender D, Maller J, Sklar P, de Bakker PIW, Daly MJ et al. . 2007. PLINK: a tool set for whole-genome association and population-based linkage analyses. Am J Hum Genet 81(3):559–575.

Quast C, Pruesse E, Yilmaz P, Gerken J, Schweer T, Yarza P, Peplies J, and Glöckner FO. 2013. The SILVA ribosomal RNA gene database project: improved data processing and web-based tools. Nucleic Acids Res 41(Database issue):D590–D596.

Raimondi S, Amaretti A, Gozzoli C, Simone M, Righini L, Candeliere F, Brun P, Ardizzoni A, Colombari B, Paulone S et al. . 2019. Longitudinal Survey of Fungi in the Human Gut: ITS Profiling, Phenotyping, and Colonization. Front Microbiol 10:1575.

Rivera WL. 2008. Phylogenetic analysis of Blastocystis isolates from animal and human hosts in the Philippines. Veterinary Parasitology 156(3):178–182.

Rothschild D, Weissbrod O, Barkan E, Kurilshikov A, Korem T, Zeevi D, Costea PI, Godneva A, Kalka IN, Bar N et al. . 2018. Environment dominates over host genetics in shaping human gut microbiota. Nature 555(7695):210–215.

Schuster FL, and Ramirez-Avila L. 2008. Current World Status of *Balantidium coli*. Clinical Microbiology Reviews 21(4):626.

Schwarz H, Tuckwell J, and Lotz M. 1993. A receptor induced by lymphocyte activation (ILA): a new member of the human nerve-growth-factor/tumor-necrosis-factor receptor family. Gene 134(2):295–298.

Shannon C. 1984. A mathematical theory of communication. .

Shieban F. 1971. Studies on Intestinal Protozoa of Domestic Pigs in the Teheran Area of Iran**This project is supported in part by funds for the Endemic Diseases Research project of plan organization and of the Department of Health Science, Teheran University School of Medicine. British Veterinary Journal 127(3):iii–v.

Summers KL, Frey JF, Ramsay TG, and Arfken AM. 2019. The piglet mycobiome during the weaning transition: a pilot study1. Journal of Animal Science 97(7):2889–2900.

Tito RY, Chaffron S, Caenepeel C, Lima-Mendez G, Wang J, Vieira-Silva S, Falony G, Hildebrand F, Darzi Y, Rymenans L et al. . 2019. Population-level analysis of Blastocystis subtype prevalence and variation in the human gut microbiota. Gut 68(7):1180–1189.

Underhill DM, and Iliev ID. 2014. The mycobiota: interactions between commensal fungi and the host immune system. Nat Rev Immunol 14(6):405–416.

Wang J, Chen L, Zhao N, Xu X, Xu Y, and Zhu B. 2018. Of genes and microbes: solving the intricacies in host genomes. Protein & Cell 9(5):446–461.

Ward-Kavanagh LK, Lin WW, Šedý JR, and Ware CF. 2016. The TNF Receptor Superfamily in Co-stimulating and Co-inhibitory Responses. Immunity 44(5):1005–1019.

White JK, Nielsen JL, and Madsen AM. 2019. Microbial species and biodiversity in settling dust within and between pig farms. Environmental Research 171:558–567.

White TJ, Bruns T, Lee S, and Taylor J. 1990. PCR Protocols amplification and direct sequencing of fungal ribosomal RNA genes for phylogenetics: Elsevier.

Whittaker RH. 1972. Evolution and Measurement of Species Diversity. Taxon 21(2/3):213–251.

Willinger T, and Flavell RA. 2012. Canonical autophagy dependent on the class III phosphoinositide-3 kinase Vps34 is required for naive T-cell homeostasis. Proc Natl Acad Sci U S A 109(22):8670–8675.

Wylezich C, Belka A, Hanke D, Beer M, Blome S, and Höper D. 2019. Metagenomics for broad and improved parasite detection: a proof-of-concept study using swine faecal samples. International Journal for Parasitology 49(10):769–777.

Xiao L, Estellé J, Kiilerich P, Ramayo-Caldas Y, Xia Z, Feng Q, Liang S, Pedersen AØ, Kjeldsen NJ, Liu C et al. . 2016. A reference gene catalogue of the pig gut microbiome. Nature Microbiology 1(12):16161.

Yang J, Lee SH, Goddard ME, and Visscher PM. 2011. GCTA: a tool for genome-wide complex trait analysis. Am J Hum Genet 88(1):76–82.

Zakrzewski M, Simms LA, Brown A, Appleyard M, Irwin J, Waddell N, and Radford-Smith GL. 2018. IL23R-Protective Coding Variant Promotes Beneficial Bacteria and Diversity in the Ileal Microbiome in Healthy Individuals Without Inflammatory Bowel Disease. Journal of Crohn’s and Colitis 13(4):451–461.

Zhao S, Xia J, Wu X, Zhang L, Wang P, Wang H, Li H, Wang X, Chen Y, Agnetti J et al. . 2018. Deficiency in class III PI3-kinase confers postnatal lethality with IBD-like features in zebrafish. Nature Communications 9(1):2639.

Zilber-Rosenberg I, and Rosenberg E. 2008. Role of microorganisms in the evolution of animals and plants: the hologenome theory of evolution. FEMS Microbiology Reviews 32(5):723–735.

