## Supplementary figure for "Gut eukaryotic communities in pigs: diversity, composition and host genetics contribution"

**Supplementary figure 1**. Sample distribution of fungi (A) and protist (B) communities. Diversity indexes of fungi (C) and protist (D) communities. Blue color represents the finishing pigs (experimental farm) and red the weaned piglets (commercial farm).

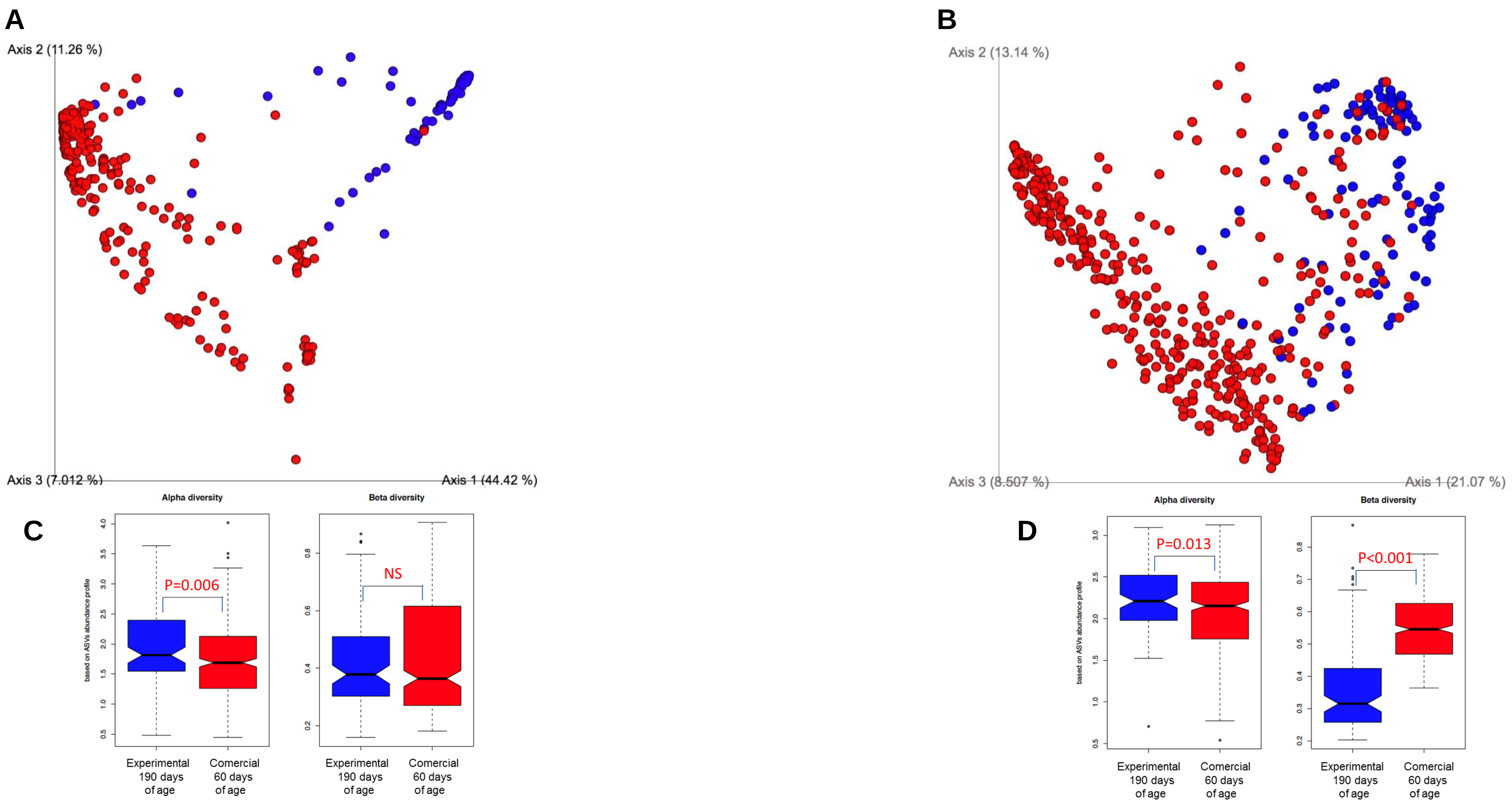
